# Peering Inside the Black Box: Explainable AI to Interpret Advanced Computer Vision Fungal Pathogen Prediction

**DOI:** 10.1101/2025.06.27.662051

**Authors:** Joshua D. Guthrie, Shamanth A. Shankarnarayan, Daniel A. Charlebois

## Abstract

Antimicrobial resistance is a growing concern, with pathogenic fungi making a substantial contribution to untreatable life-threatening infections across the globe. Artificial intelligence is increasingly used in microbiology and antimicrobial resistance research, with promise to improve clinical diagnostics and infectious disease treatments. However, how AI models make predictions remains largely unknown, which is a major hurdle for human trust and regulatory approval. We trained convolutional neural networks (DenseNet121 and InceptionV3) and vision transformers (Swin Transformer-Tiny and Vision Transformer-Base 16) to quickly and accurately identify human fungal pathogens from microscopy images. Using explainable AI (Occlusion Sensitivity and Grad-CAM), we identified biologically relevant features (organelle, cell interior, cell wall, budding patterns/scars, and optical patterns) and irrelevant image features (background artifacts) high-performance computer vision models used to make predictions. These findings advance our understanding of how computer vision models make predictions on microbial pathogens and are anticipated to have profound implications for AI-based diagnostics.

## 1 Introduction

Globally, 6.5 million fungal infections are estimated to occur annually and are associated with 3.8 million deaths, of which 2.5 million deaths are directly attributed to these infections^1^. *Candida* bloodstream infection (candidemia) and invasive candidiasis result in nearly 1 million deaths annually^1^. Many fungal infections are underreported and underdiagnosed, and treatment is restricted to a few classes of antifungal drugs^2^. Candidemia is the fourth most common infectious disease in intensive care units and occurs predominantly in hospitalized patients with underlying risk factors; however, candidemia is not limited to patients who are immune-compromised and/or critically ill^3^. The emergence of multiple new human fungal pathogens each year^2^ and increasing resistance to antifungal drugs has escalated the demands on healthcare facilities, leading to higher economic burden and mortality^4^. *Candida albicans* is the leading cause of candidemia in many geographical locations around the world. However, non-albicans species such as *Nakseomyces glabratus* (previously *Candida glabrata*), *Pichia kudriavzevii* (previously *C. krusei*), *Candida parapsilosis*, and *Candida tropicalis* contribute up to 95% of the disease burden^3,5^. The multidrug-resistant pathogenic yeast *Candidozyma auris* (previously *Candida auris*) has appeared in over 60 countries over the last decade, with hospital-acquired infections and outbreaks being a major concern^6^.

Multiple challenges exist to identify pathogenic yeast species. The blood culture method is the gold standard for identifying yeasts in candidemia cases^7^. Alternatively, non-culture-based methods, such as identifying serum biomarkers (*β* -(1,3)-D-glucan) and nucleic acids (nucleic acid amplification tests)^8^, are also employed. However, these methods are limited due to low sensitivity, long turnaround times, and sophisticated instrumentation, specialized laboratory space, and highly-trained personnel requirements^7^. To overcome these challenges, several new detection methods are being developed for faster detection of yeast species directly from blood samples, as well as yeasts grown on agar plates^7^. These detection methods include the use of chrome agar^9^, matrix-assisted laser desorption/ionization-time of flight mass spectroscopy (MALDI TOF MS)^10^, DNA sequencing^11^, T2 magnetic resonance^12^, volatile organic compounds^13^, Fourier transform infrared with attenuated total reflectance^14^, imperfect match probes^15^, recombinase polymerase amplification^16^, high resolution melting curves^17^, Raman spectroscopy^18^, and laser-induced breakdown spectroscopy^19^. However, many challenges exist, such as integrating these methods into routine diagnostic workflows, species-level misidentification, and limited access in low-and middle-income countries^1^.

The use of artificial intelligence (AI) and machine learning (ML) has seen remarkable growth in recent years, driven by advances in model architectures, training algorithms, computational hardware, open-source AI frameworks (e.g., PyTorch^20^), and the availability of vast amounts of training data^21^. Artificial intelligence and ML approaches are increasingly being applied in scientific research to model complex data^21^. For instance, AlphaFold can determine the 3D structures of proteins and their bimolecular interactions with accuracies comparable to experiments, and is accelerating drug discovery^22^. Recent studies have investigated ML methods in the diagnosis and treatment of infections^23^, and to discover antibiotics^24^. The field of computer vision, which applies AI to analyze image data, has made substantial progress from advances in artificial neural networks and attention-based transformer models^25–27^. Computer vision models based on convolutional neural network (CNN) and vision transformer architectures have achieved state-of-the-art performance on gold-standard image classification tasks^25–28^, such as the ImageNet competition^26^. These models have been applied in biological and medical image analysis^29^, achieving performance comparable to current diagnostic tools and human professionals^30^. They have also been applied in recent studies for the classification of yeast species in microscopy images^31–33^. Furthermore, advances in computer vision segmentation models have enabled advanced automated image processing^34^, finding various applications in cell microscopy image analysis^35–38^.

Deep learning model architectures have been developed for computer vision applications. Deep learning methods have been applied in the context of yeast pathogens, including segmenting microscopy images^37,39^and inner organelles^40^, tracking cellular growth^41^, and measuring cell dimensions^42^. InceptionV3^43^, a deep CNN architecture with 48 layers and “inception modules” that apply convolutions at various filter sizes to capture features at multiple scales, was previously combined with transfer learning to classify *Candida* species^33^. Specifically, the InceptionV3-based model was able to classify four pathogenic yeast species (*C. albicans, C. auris, N. glabratus*, and *C. haemulonii*) from brightfield microscopy images with an overall accuracy of 78%. DenseNet121, ViT, and Swin Transformer models have not been used for yeast species classification and provide improvements over InceptionV3 and other computer vision architectures^25,44–46^. DenseNet121 is a deep CNN architecture with 121 layers that features a dense connectivity pattern that connects each layer in the network to all preceding layers^44^. The DenseNet architecture was designed to improve gradient calculations and promote the reuse of features, leading to better efficiency through faster optimization and smaller network and dataset size requirements to reach high performance^44^. Vision transformers are relatively new models that depart from the CNN architecture by adapting the transformer architecture of large language models to computer vision tasks, using self-attention mechanisms to better learn global features in images^45^, resulting in comparable or better results than CNNs on standard image classification metrics^45^. However, vision transformers require more data than CNNs to reach comparable performance^25^. Swin (Shifted Window) transformers are a modification of the vision transformer architecture that addresses this limitation through a hierarchical representation of feature maps and the use of shifting window attention, improving local feature extraction capabilities while reducing computational complexity^46^.

The “black box” nature of AI models often renders their predictions difficult, or even impossible, to understand. This limits the application of AI in fields where model interpretation and human trust are of particular importance, such as scientific research and medical diagnostics^47^. The emerging field of explainable AI (XAI) aims to elucidate how AI/ML models make predictions^48^, with an emphasis on computer vision^49^. Although complete explainability of computer vision models has yet to be achieved, XAI methods can identify image features that are most important for model predictions^49^. Understanding the decision-making process underlying AI models will enable scientific and medical fields to adopt AI-driven tools with greater confidence. Explainable AI methods investigate predictions made by computer vision models, such as Grad-CAM (Gradient Class Activation Mapping)^50^ and Occlusion Sensitivity^51^. Grad-CAM uses information flowing into the final convolutional layer of a CNN to produce a coarse-grained visualization of areas in an image that were most important for a given prediction^50^. This makes Grad-CAM suitable for determining if the model places high prediction importance on a high-level object within an image, but not for determining specific predictive features. Occlusion Sensitivity is a model-agnostic perturbation method that measures the change in prediction output of a model when occluding specific regions of the input image, with large changes in model output indicating prediction importance of the occluded region^51^. Occlusion Sensitivity is capable of producing fine-grained importance visualizations to identify specific features that lead to model predictions^51^.

In this study, we develop high-performance computer vision models (InceptionV3^43^, DenseNet121^44^, Vision Transformer (ViT)-Base 16)^45^, and Swin Transformer-Tiny^46^) to quickly and accurately identify yeast pathogens from microscopy images. We overcome current limitations in computer vision pathogen identification by applying XAI to interpret the predictions made by these advanced computer vision classification models. We use segmentation models to automate data processing, enabling the generation of large datasets for model training and evaluation, while increasing data processing efficiency, eliminating a major bottleneck of previous work^31–33^. This increase in training data and improvements in model development, particularly the use of state-of-the-art model architectures and full model fine-tuning, leads to substantial improvements in model performance that scales with classification complexity through the inclusion of additional pathogenic yeast species (e.g., *P. kudriavzevii, C. parapsilosis*, and *C. tropicalis*). The use of XAI methods to interpret computer vision predictions enables model verification, determines biologically relevant and irrelevant features used for species predictions, identifies areas for model improvement, and enhances trust in the application of AI models for pathogen identification.

## 2 Materials and Methods

### 2.1 Sample Preparation, Imaging, and Data Processing

Clinical isolates of *C. albicans, C. auris*, and *N. glabratus* were obtained from the Alberta Precision Laboratory -Public Health Laboratory. the The American Type Culture Collection (ATCC) provided the strains for *C. parapsilosis* ATCC-22019, *P. kudriavzevii* ATCC-6258, *C. haemulonii* ATCC-22991, and *C. tropicalis* ATCC-750. All strains were preserved in 25% glycerol at -80°C until further use. The clinical and standard strains were revived by culturing from frozen stock in Sabouraud glucose agar (SDA) plates (Millipore, Darmstadt, Germany, 1.05438.0500) and incubated at 35°C for 48 h. Fresh subcultures were grown on SDA agar plates and incubated at 35°C for 24 h. Isolated colonies of different pathogenic yeast species grown in SDA medium for 24 h were used to prepare wet mount slides (Fisherbrand, Pittsburg, USA, 22-034486) using sterile distilled water, overlayed with 22 mm coverslips (Fisherbrand, Pittsburg, USA). The raw images for each species were captured using the EVOS M7000 imaging system (Invitrogen, Thermo Fisher Scientific, Massachusetts, USA, AMF7000) with an 100x oil immersion objective l. Coarse and fine adjustments were made to focus the cell on a single plane prior to capturing images. The brightness of the trans-illumination was adjusted to 0.033 before image capture.

Images of individual cells were cropped from the raw microscope images to build training, validation, and test datasets^33^. This process was automated using a CNN-based yeast cell segmentation model (yeaZ) for brightfield images^37^. Applying yeaZ produced segmentation masks for each raw microscope image, which contained the boundary coordinates of the identified yeast cells. To ensure each cell was fully contained within its segmentation border, the boundary coordinates were expanded by a factor of 0.15 using morphological dilation (implemented with scikit-image^52^). The expanded boundary coordinates were used to calculate square (1:1 aspect ratio) bounding boxes around each identified cell, which were then used to crop individual cell images with a padding of 10 pixels. For each crop, pixels that were outside of the expanded cell segmentation boundary coordinates were set to 0 (black) to remove the background around each cell, reducing irrelevant image features. Due to intrinsic errors of the segmentation model and image data, a large number of low-quality images (e.g. slide artifacts and poorly segmented cells) were cropped using this process. To automate the removal of these images, crops that were smaller than 50×50 pixels were excluded (as these were generally found to be slide artifacts), and feature extraction with ResNet18^53^ and k-means clustering^54^ with k=2 were used to separate the remaining high-quality crops from low-quality crops^55^, which we found to produce consistent results for our datasets. This process was applied to the raw microscope images collected for training, validation, and testing. To ensure dataset balance and avoid biases during the training and validation process^56^, the training and validation sets were randomly downsampled to match the species with the lowest number of crops in each set (each species had a different number of individual cell images due to the random distribution of cells across the raw microscopy images). The test set was randomly downsampled to match the size of the validation set to maintain consistency when comparing model validation and test results.

### 2.2 Image Dataset Collection

A total of 900-1,200 raw microscope images for model training, 100-300 for validation, and 100-250 for testing were collected for each species from different subcultures. The data processing system described in Section 2.1 was applied to the raw images, resulting in 32,809 training, 4,214 validation, and 4,214 test images after k-means to remove low-quality crops and randomly downsampling to maintain balanced datasets. Although k-means removed most low-quality crops, some still appeared in the final datasets, particularly for *C. auris* and *P. kudriavzeviii*. This was likely due to the smaller cell size and other morphological differences compared to the yeast species that the segmentation model was trained on (*S. cerevisiae*)^37^, which may have caused challenges during segmentation and clustering. The outlier rates in the test set were quantified by visually inspecting the images and counting the number of outliers for each species, showing that 2.4% of the overall test set consisted of low-quality crops that slipped through detection (705 out of 7 × 4, 214 = 29, 498 images). Most species had an outlier rate of 1% or less, while *C. auris* and *P. kudriavzevii* had relatively high rates of 9.1% (387 outliers out of 4,214 images) and 4.8% (204 outliers out of 4,214 images), respectively. To ensure that these outliers (artifacts and poor crops) did not bias our test results, we manually removed them from the test datasets. Model test performance without manual outlier removal was comparable to test performance with manual outlier removal (Supplementary Section 3, Supplementary Figure 5, and Supplementary Table 4). The number of images for each species was then randomly down-sampled to maintain a balanced test set, resulting in 3,827 individual cell images per species. The training, validation, and test raw microscopy images and individual cell crops after each data processing step for each species are summarized in Supplementary Table 1. Pixel and image size distributions for the final training, validation, and test datasets are shown in Supplementary Figures 1, 2, and 3, respectively.

### 2.3 Computer Vision Model Training and Evaluation

Transfer learning and fine-tuning were used to train the computer vision models^57^ to classify pathogenic yeasts. The pre-trained base network acts as a generalized model of image classification tasks, while the added network learns features specific to the new training data to enable pathogen classification. To adapt the models to microscopy images of pathogenic yeast species, fine-tuning was applied to the full networks (the base model architecture with a randomly initialized 7-class classification head) by initializing the base model with ImageNetV1 weights and then training the entire network using the microscopy image training dataset. This process adjusted the pre-trained weights to enable the network to learn the visual characteristics specific to the pathogenic yeast species, leveraging the advanced network architectures of the base models and their pre-trained weights to improve the classification performance of the models.

Four computer vision architectures were considered for the base model, implemented using PyTorch^20^: InceptionV3^43^, DenseNet121^44^, a base 16 Vision Transformer (ViT)^45^, and a Swin Transformer-Tiny^46^. Image transformations (resizing, rescaling, and standardization) specific to each pre-trained model (based on the ImageNetV1 dataset^58^) were applied to all images before they were used for training, validation, and testing. Each model was trained on the training set using an AdamW optimizer^59^ to minimize a cross-entropy loss function^60^. To ensure consistent model comparisons, the same hyperparameters were used to train each model, with a learning rate of 3e-5, an AdamW weight decay of 0.1, and a batch size of 32. The validation set was used to evaluate model performance (overall accuracy and loss) after each iteration (epoch) of the training process. Early stopping^61^ (with a patience of 5) was used to stop the training process once the validation loss no longer decreased, with the model weights from the epoch with the best validation performance (lowest validation loss) saved. After the training and validation process was completed, the models were applied to the test set to evaluate their unbiased generalization performance. Standard classification metrics were used to quantify model performance, including the overall model accuracy and class-specific precision, recall, and *F*_1_-score, and their averages over all classes. Precision is defined as the ratio between the true positive (TP) and all positive predictions (TP plus false positives) for a given class. Recall and class accuracy are used interchangeably in this study and defined as the correct predictions (TP) divided by the total number of samples (TP plus false negatives) for a given class. The *F*_1_-score, defined as the harmonic mean between precision and recall, was used to evaluate the balance between these metrics. Confusion matrices were used to visualize model performance and identify classification trends. The classification metrics, confusion matrices, and principal component analysis (PCA) were determined using scikit-learn^62^.

To investigate the feature space of the images for each species, we performed a PCA^63^ on the test set images after using our trained model as a feature extractor. PCA reduced the dimensionality of the feature representations learned by our trained model to visualize their principal 2D and 3D components. This enabled us to identify differences and similarities between the representations of each species and to better understand common misidentification patterns that occurred ruing model evaluation by showing species that overlap in the feature space.

### 2.4 Explainable AI for Model Verification and Prediction Interpretation

We applied XAI methods to determine if the high-performance computer vision models were learning relevant features (e.g., cellular characteristics such as cell wall and organelles) or irrelevant features (e.g., background artifacts). Additionally, we used XAI to provide additional verification to model predictions and identify features that were most important for species predictions. We chose to use Grad-CAM^50^ and Occlusion Sensitivity^51^ for our image classification models. Grad-CAM was chosen because it has been extensively tested^64^ and used for computer vision model interpretation^65^. Occlusion Sensitivity was selected because it does not rely on specific model architectures^51^ and is capable of producing more fine-grained feature importance visualizations than Grad-CAM. Grad-CAM and Occlusion Sensitivity, implemented using the PyTorch-based library Captum^66^, produce normalized attribution maps (values ranging from 0 to 1) that highlight regions of the input image with high importance for a given model prediction. Different sets of test predictions (correct and incorrect) made by the model were investigated by generating Grad-CAM and Occlusion Sensitivity attribution maps for a large number of input images and visually inspecting them to identify image features with the highest attribution (equal to 1). The results of the visual inspection were then consolidated by calculating the proportion of images in the set that contained high attribution on each of the features.

First, we used Grad-CAM to distinguish whether there was high attribution inside the cell, outside the cell, or partly inside and outside the cell in each image. This resulted in three measurable binary variables (high attribution yes/1 or no/0 for each case) for each attribution map. Next, Occlusion Sensitivity (with a small non-overlapping 2×2 pixel sliding window) was used to produce fine-grained attribution maps to identify specific image features contributing to model predictions. We identified nine visually distinguishable high-level features that could be present in the input images: internal organelles (nucleus and/or vacuoles), the cell interior (not on a specific organelle), the inner optical pattern, the cell wall, the outer optical pattern, budding patterns/scars, partially cropped cells outside of the main cell, the slide background (between the cell and the background removal border), and the black background. This resulted in nine measurable binary variables for each Occlusion Sensitivity attribution map (high attribution yes/1 or no/0 for each feature), with six of the features being biologically relevant (organelle, cell interior, cell wall, budding pattern/scar, inner and outer optical patterns) and three of these features being irrelevant (other cells in the image, slide background, and black background). As more than one feature had high attribution in the majority of cases, each attribution map generally had more than one high-importance feature.

The data collection/processing, model training and evaluation, and XAI workflow are shown in Figure 1.

**Figure 1.**
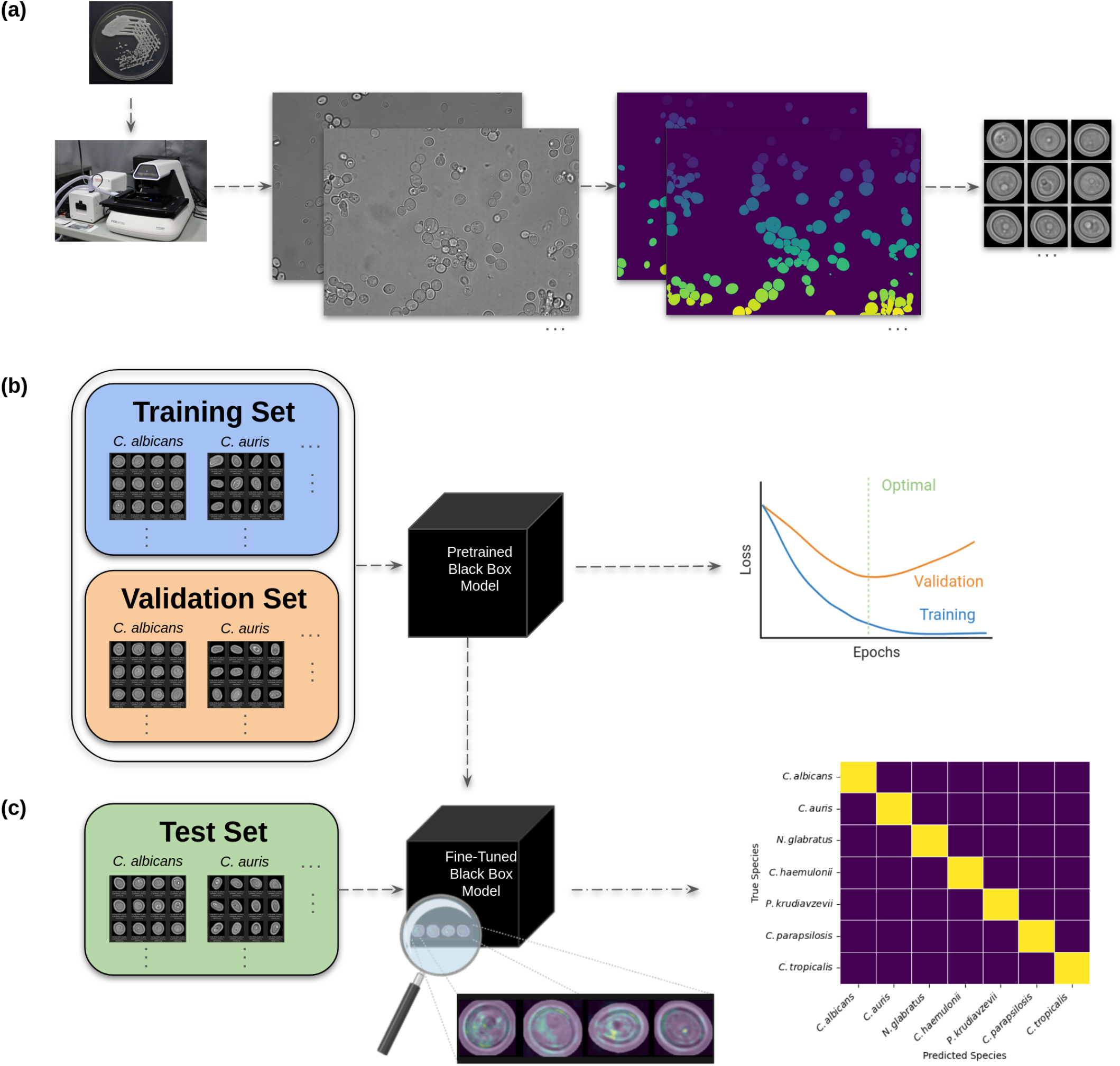
Workflow for training and evaluating the advanced computer vision pathogenic yeast species classification models. (a) Raw microscope images for each pathogenic species were collected in three independent groups to create training, validation, and test datasets. Automated data processing was used to crop images of individual cells from the raw microscopy images. (b) The training data set was used to train “black-box” computer vision models, with model performance being evaluated on the validation set after each epoch to optimize hyperparameters. (c) The test data set was used to evaluate model generalization performance, visualized using confusion matrices. Explainable AI was used to interpret model predictions and to identify biologically relevant and irrelevant image features with high prediction importance. Parts of this figure were created using Bio-Render.com.

## 3 Results and Discussion

### 3.1 Advanced Computer Vision Pathogenic Yeast Classification

We developed high-performance computer vision models to classify seven pathogenic yeast species (Section 2.3). Each computer vision model produced comparable overall test results, with DenseNet121 performing marginally better than the other models (Figure 2). Optimal performance (lowest validation loss) for the DenseNet121 model was obtained after epoch 5 with a training loss, training accuracy, validation loss, and validation accuracy of 0.148, 95%, 0.367, and 88%, respectively (Figure 3a). The corresponding validation confusion matrix (Figure 3b) and the validation performance metrics (Supplementary Table 2a) were also determined. Applying DenseNet121 to the test set resulted in an overall test accuracy of 87% with strong classification performance across all species, correctly classifying *C. albicans, C. auris, N. glabratus, C. haemulonii, P. kudriavzevii, C. parapsilosis*, and *C. tropicalis* in 89%, 86%, 85%, 76%, 89%, 97%, and 82% of cases, respectively (Figure 4 and Supplementary Table 2b). The test performance was generally balanced between precision and recall across the species, with *F*_1_ scores ranging from 0.83 to 0.92 with an average value of 0.87 (Supplementary Table 2b). The most substantial differences between precision and recall were for *C. parapsilosis* (0.73 precision, 0.97 recall) and *C. haemulonii* (0.90 precision, 0.76 recall), with specific predictions contributing to these imbalances (Figure 4). The relatively low precision of *C. parapsilosis* arose due to other species, primarily *N. glabratus, C. haemulonii*, and *C. auris*, being misidentified as *C. parapsilosis*, while the low recall of *C. haemulonii* was due to it being misidentified as other species, primarily *C. parapsilosis, C. albicans*, and *N. glabratus*. The test set confusion matrices and performance metrics for the other three computer vision models were also obtained (Supplementary Section 2, Supplementary Figure 4, and Supplementary Table 3). All models produced similar test results and classification patterns (Figure 4 and Supplementary Figure 4), with DenseNet121 producing the best results for *N. glabratus* and *C. haemulonii* (Figure 2). To investigate how outliers affect these results when using a fully automated test set without manual outlier removal, we applied the DenseNet121 model to a test set containing outliers (Supplementary Section 3, Supplementary Figure 5, and Supplementary Table 4). The inclusion of outliers resulted in a 1% decrease in overall test accuracy, with a minor effect on performance for all species except *C. auris* and *P. krudiavzevii*, which respectively resulted in a 4% decrease in recall and a 4% decrease in precision.

**Figure 2.**
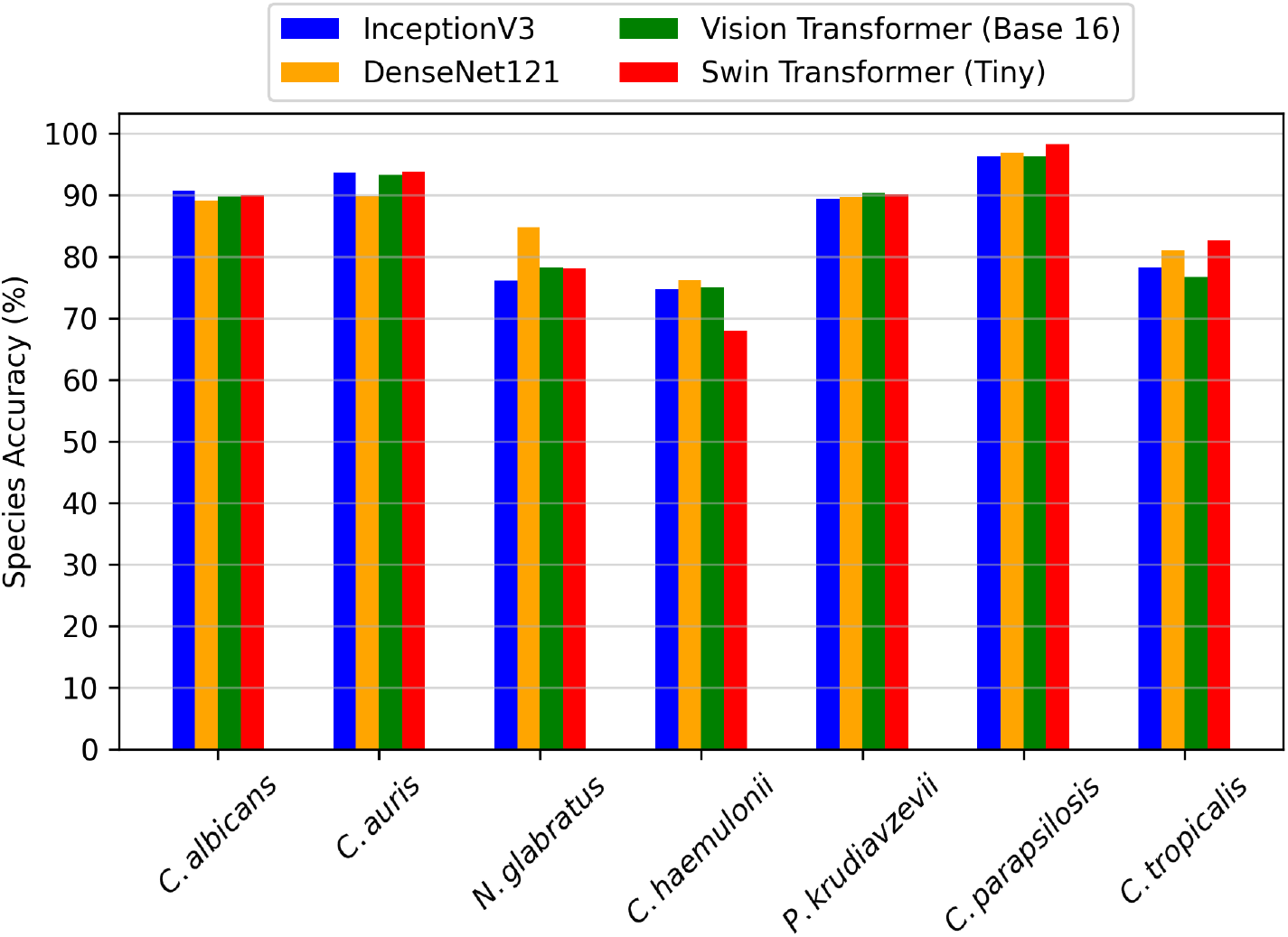
Pathogenic yeast species classification accuracies for advanced computer vision models. DenseNet121, Swin Transformer (Tiny), Vision Transformer (Base 16), and InceptionV3 models had overall test accuracies of 87%, 86%, 86%, and 86%, respectively.

**Figure 3.**
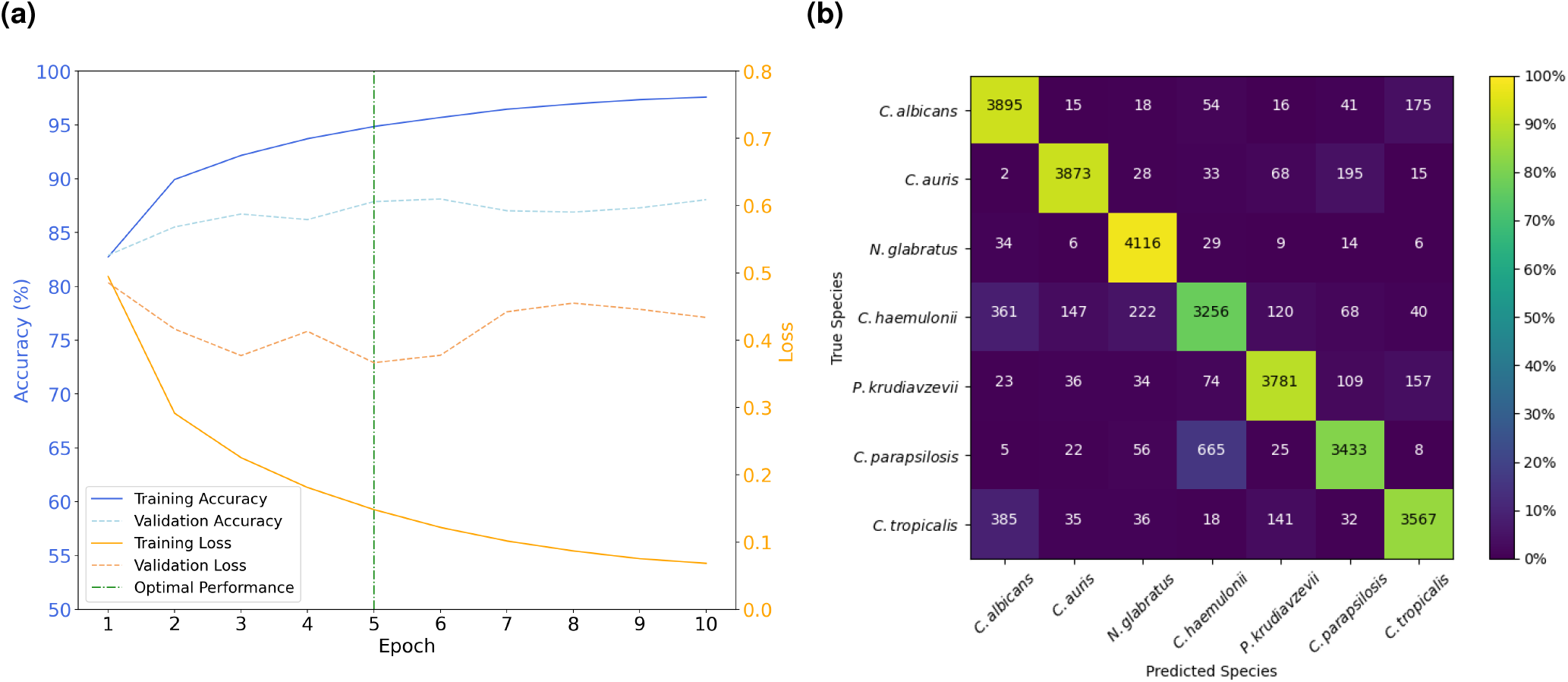
Training and validation performance of the DenseNet121 pathogen classification model. **(a)** Training and validation accuracy and loss curves. The model weights from the optimal epoch (epoch 5) were saved, which had a validation and training accuracy of 95% and 88% and loss of 0.148 and 0.367, respectively. **(b)** Confusion matrix for the DenseNet121 model applied to the validation set. The heatmap shows the percentage of samples in each cell relative to the validation dataset size of each class (4214 images).

**Figure 4.**
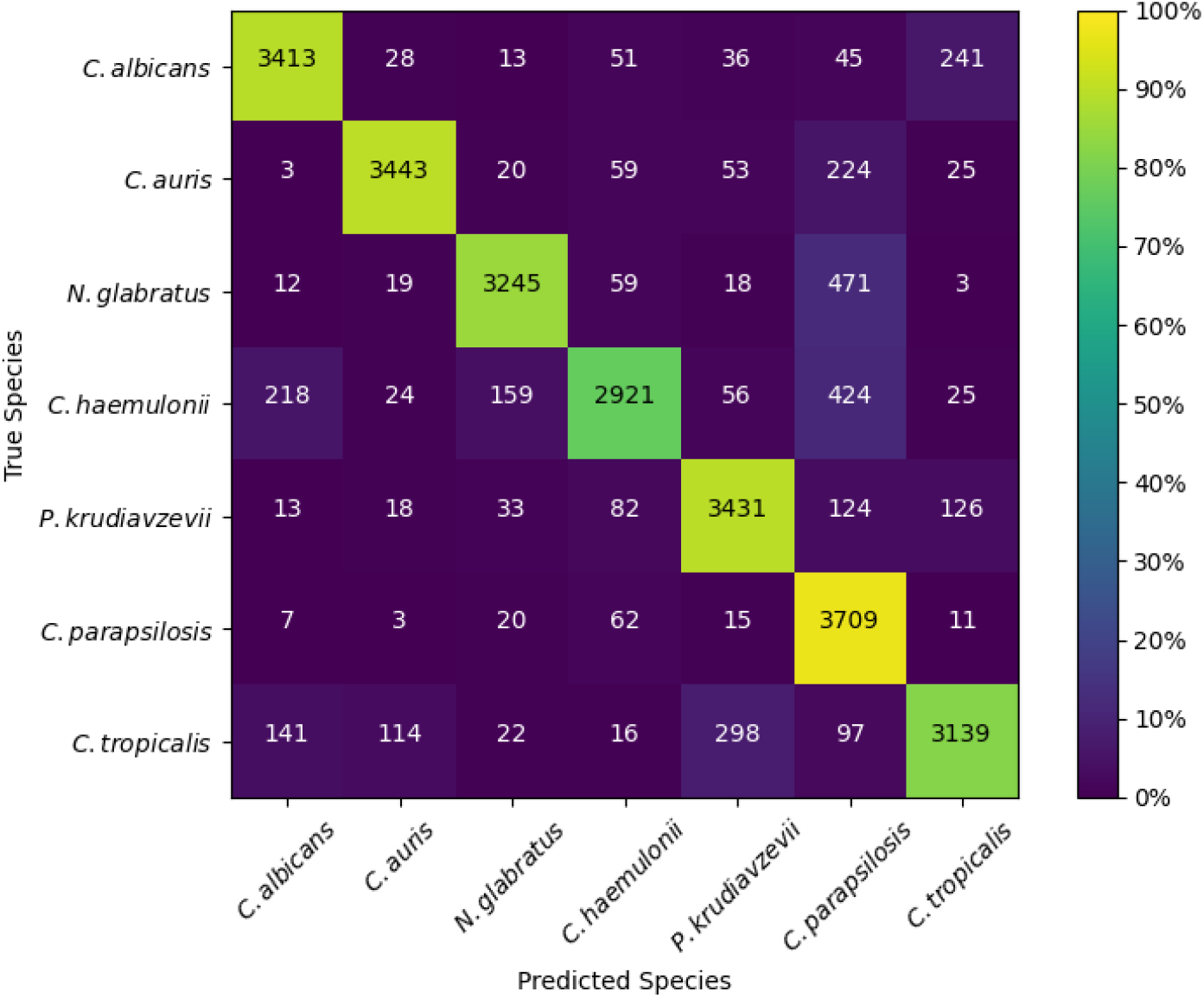
Confusion matrix of test set predictions made by the DenseNet121 pathogen classification model. The heatmap shows the percentage of samples in each cell relative to the test dataset size of each class (3827 images).

To investigate the feature characteristics of the test set for each species used by our DenseNet121 model to make predictions, and to identify reasons for model confusion, we performed a PCA. The PCA revealed that species that were confused by the DenseNet121 model have feature spaces with similar shapes and extensive overlap, for example *C. albicans* and *C. tropicalis* (Figure 5 and Supplementary Video 1). Furthermore, species that the model could distinguish between had minimal overlap in the PCA, including *C. albicans* and *C. auris* (Figure 5). In some cases, the model confused one species for another without the reverse being true. For instance *N. glabratus* was often misidentified as *C. parapsilosis*, but not vice versa (Figure 4. The PCA results indicate that these scenarios occur when a species with a spread-out feature space, such as *N. glabratus*, overlaps with a species that has a more concentrated feature space, such as *C. parapsilosis* (Figure 5). This suggests that our DenseNet121 model is biased toward species with more concentrated feature spaces because a lower variance in visible features is easier for the model to learn. This likely occurs due to model confusion when an image of a species with more variability in visible features contains features similar to another species, resulting in inaccurate species predictions.

**Figure 5.**
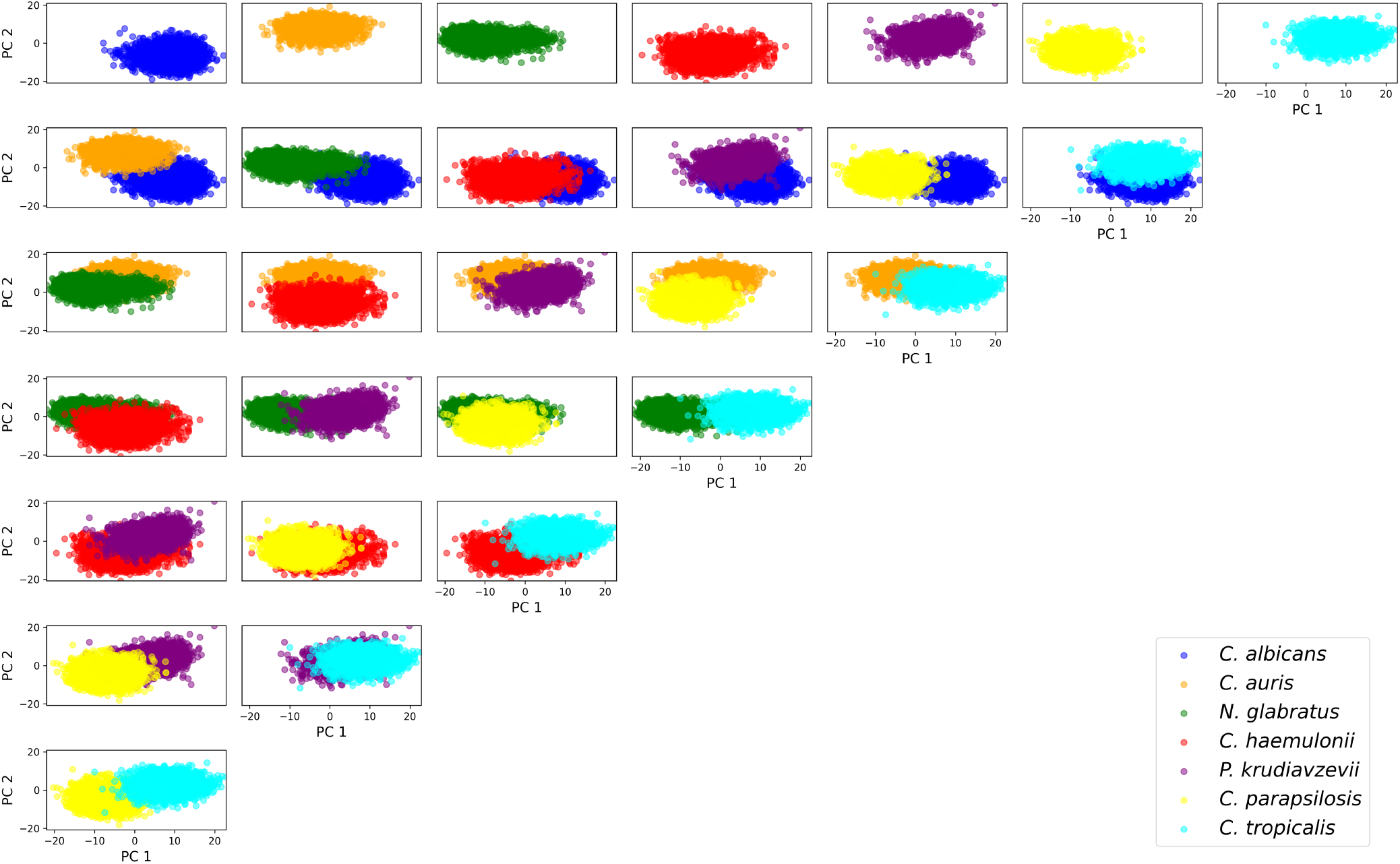
A 2-dimensional principal component analysis (PCA) of test set images for each pathogenic yeast species using the trained DenseNet121 model as a feature extractor. The first row shows the PCA for individual species, followed by rows showing comparisons of species pairs. A 3-dimensional PCA is provided in Supplementary Video 1.

Next, we trained, validated, and tested the high-performance computer vision models using our new approach on four yeast species previously considered to compare with an existing machine learning-based pathogenic yeast classifier^33^. Performance was similar across each of our model architectures, with the Vision Transformer producing the best test results, having an overall accuracy of 92% and correctly classifying *C. albicans, C. auris, N. glabratus*, and *C. haemulonii* in 95%, 97%, 96%, and 82% of cases, respectively (Supplementary Section 4 and Supplementary Figure 6). This is a substantial improvement over the previously published test results of the older model, which had an overall accuracy of 78% and correctly classified *C. albicans, C. auris, N. glabratus*, and *C. haemulonii* in 97%, 73%, 69%, and 73% of cases, respectively, highlighting the improvements we have made in data collection, processing, and model development. The model architecture test set comparisons, the Vision Transformer confusion matrix, and a species accuracy comparison to previous work^33^ are shown in Supplementary Figure 6.

**Figure 6.**
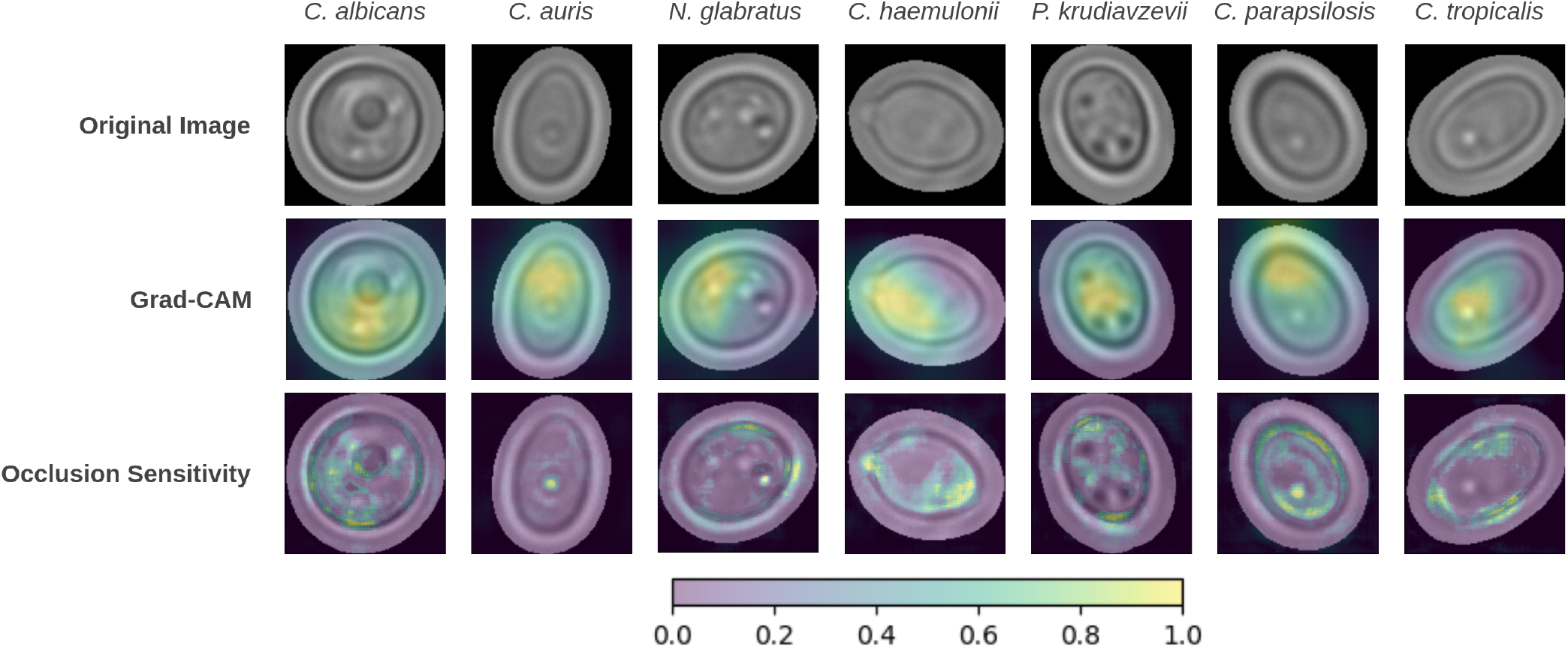
Representative cell images with overlayed Grad-CAM^50^ and Occlusion Sensitivity^51^ attribution maps for true positive test set predictions made by the DenseNet121 pathogen classification model. The colour bar shows the normalized attribution values.

### 3.2 Explainable AI to Interpret Computer Vision Model Predictions

The size and budding patterns of pathogenic yeast cells were previously hypothesized to contribute correct species classification by computer vision models^33^. However, this hypothesis could not be verified due to the black-box nature of AI models^47^. Here, we used XAI, namely Grad-CAM^50^ and Occlusion Sensitivity^51^, to interpret the predictions of high-performance computer vision models. Specifically, we used these XAI methods to identify image features that these improved computer vision models used to accurately identify the species of pathogenic yeast. Specifically, we used Grad-CAM^50^ and Occlusion Sensitivity^51^ to generate attribution maps for three sets of one hundred randomly selected TP test set predictions for each species made by our best-performing DenseNet121 model. Additionally, we generated attribution maps for the eight most frequent DenseNet121 misidentification cases (Figure 4), namely *N. glabratus* misidentified as *C. parapsilosis, C. haemulonii* as *C. parapsilosis, C. tropicalis* as *P. kudriavzevii, C. albicans* as *C. tropicalis, C. auris* as *C. parapsilosis, C. haemulonii* as *C. albicans, C. haemulonii* as *N. glabratus*, and *C. tropicalis* as *C. albicans*, which contributed to 12%, 11%, 8%, 6%, 6%, 6%, 4%, and 4% of the test predictions for the corresponding true species, respectively. Representative Grad-CAM and Occlusion Sensitivity attribution maps are shown in Figure 6, which demonstrate the level of detail and features that each of these methods identify.

We calculated the proportion of TP predictions made by DenseNet121 with high Grad-CAM attribution inside the cell, outside of the cell, or both for each pathogenic yeast species (Figure 7a). These results showed that there was high prediction importance fully inside the cell in nearly all cases for each species, with no attribution maps indicating high importance fully outside of the cell. For the top eight misidentification cases, the proportion of images with high Grad-CAM attribution was 100% for all species (Figure 7b). These results indicate that the model places high prediction importance on the cell in each image in all correct and incorrect predictions investigated. However, the low resolution of the Grad-CAM attribution maps did not allow specific cell characteristics that contribute to model predictions to be identified. Therefore, we applied Occlusion Sensitivity to produce fine-grained attribution maps to identify specific high-importance image features (Section 2.4).

**Figure 7.**
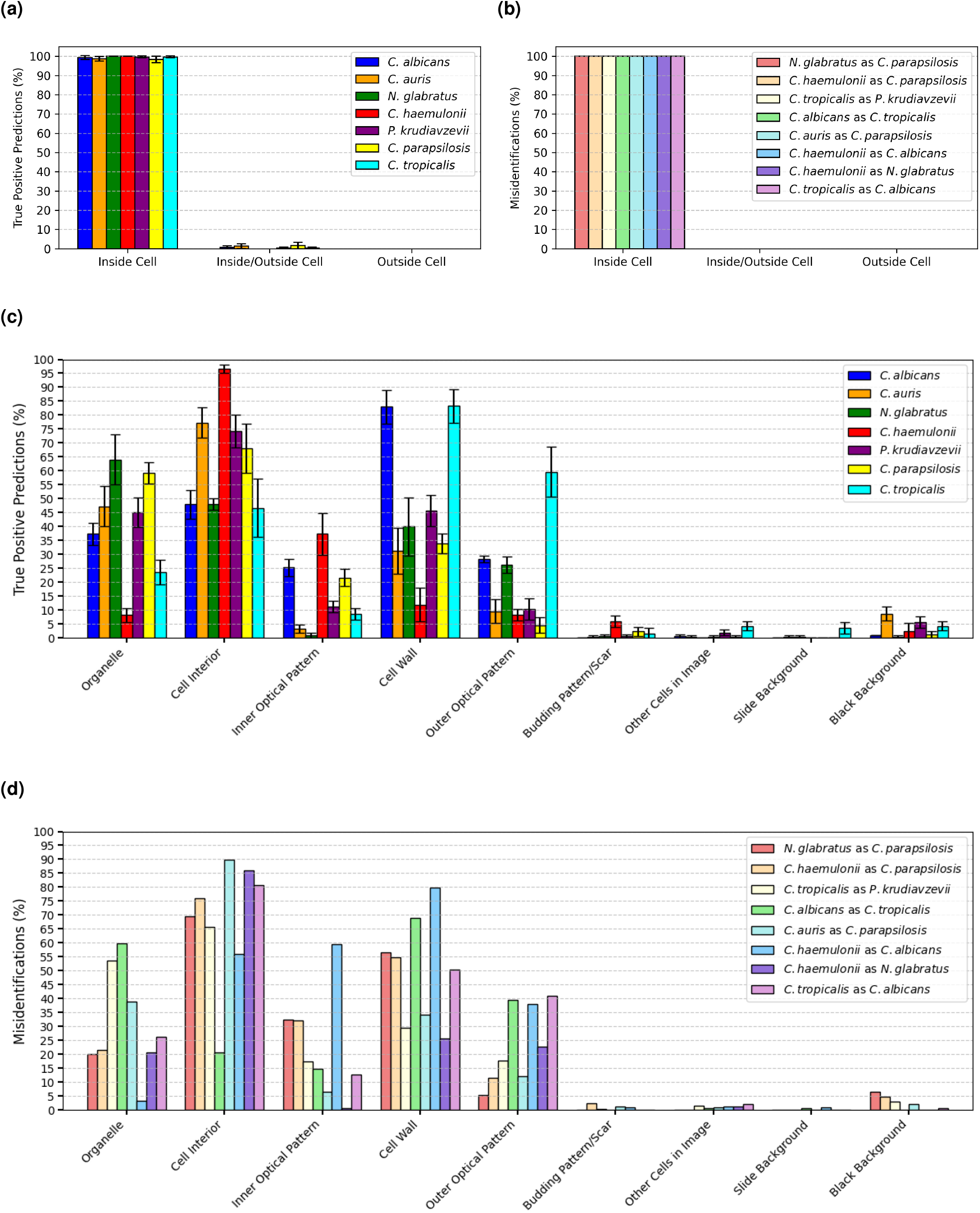
Proportions of identified features with high explainable AI attribution for test set predictions made by the DenseNet121 pathogen classification model. Grad-CAM^50^ results are shown for true positive (TP) predictions in **(a)** and for misidentification predictions in **(b)**. Occlusion Sensitivity^51^ results are shown for TP predictions in **(c)** and for misidentification predictions in **(d)**. TP results were averaged over 3 sets of 100 randomly sampled predictions from the test set for each pathogenic yeast species, with the error bars showing the standard deviation. No error bars are shown in the misidentification results, as all images for these cases were analyzed. The proportions of each case do not sum to 100%, as many images had high attribution on more than one feature.

Next, we calculated the proportion of TP predictions with high Occlusion Sensitivity attribution on each high-level feature for all species (Figure 7c). These results indicate that the DenseNet121model places high importance on biologically relevant features in the majority of TP predictions, while placing little importance on non-biologically relevant features. The most frequent high-importance feature was found to be the cell wall for TP predictions of *C. albicans* and *C. tropicalis*, the cell interior (not specific to an organelle) for *C. auris, C. haemulonii, P. kudriavzevii*, and *C. parapsilosis*, and internal organelles (nuclei and vacuoles) for *N. glabratus*. Although these were the features with the most frequent high attribution, the predictions generally had more than one feature with high attribution, and a combination of feature characteristics likely contributed to each model prediction. As for model misidentifications, we again found that the features with high Occlusion Sensitivity attribution were biologically relevant in the vast majority of cases (Figure 7d). The cell wall was the most frequent high-importance feature for *P. kudriavezevii* misidentified as *C. tropicalis, C. albicans* as *C. tropicalis*, and *C. haemulonii* as *C. albicans*, while the cell interior was the most frequent for the other five misidentification cases.

The Grad-CAM and Occlusion Sensitivity XAI methods showed that the cell in each image had the highest importance for species classification predictions made by the DenseNet121 model, indicating that the model relied primarily on biologically relevant cell features to make predictions (Figures 7). The detailed Occlusion Sensitivity attribution maps identified specific biologically relevant cell characteristics and irrelevant image features that had the highest importance for given DenseNet121 predictions (Figure 7c and 7d), indicating that the interior cell structure, cell wall, and organelles were of the highest importance for distinguishing between the species, followed by the optical patterns around the cells produced by diffraction of the microscope light source and the optical properties of the cells^67^.

## 4 Conclusion

Driven by advances in data collection and processing, model architectures, and training methods, we developed high-performance computer vision models with state-of-the-art generalization performance for human fungal pathogens. Relying only on bright-field microscopy images, this approach does not require sophisticated instruments or expertise and can be integrated into current diagnostic processes^7^. The increase in performance of our advanced computer vision models to more accurately classify a wider range of pathogenic yeasts compared to previous work^31–33^was achieved through technical improvements in microscopy image collection, data processing, and model development. Specifically, we obtained a substantially larger training dataset of individual cell images (32,809 per species) with lower time and resource costs compared to previous studies by implementing the yeaZ yeast segmentation model^37^ and the k-means algorithm^54,55^to automate raw microscope image processing. Along with full model fine-tuning, this increase in training data enabled our advanced vision models to learn nuanced morphological differences between pathogenic yeast species.

The application of XAI methods revealed that biologically relevant features are of high importance in correct and incorrect species predictions made by our advanced computer vision models. Grad-CAM^50^ was used for high-level attribution maps, whereas Occlusion Sensitivity^51^ was used to elucidate finer details. Grad-CAM showed that computer vision models place high importance on the cell in each image to make predictions, while Occlusion Sensitivity identified important features unique to each pathogenic yeast species, including the cell wall, internal organelles, cell interior structure, and optical properties. The predictions made by our best-performing model, DenseNet121, were determined by the XAI methods to be primarily based on cell morphology, enhancing transparency and trust in model predictions^47^. The XAI methods used in our study generated insights into species-specific features (e.g., optical patterns generated by cellular organelles) that are most important to distinguish yeast species and identified reasons (e.g., morphological similarities between species) for incorrect model predictions.

The similarity in performance between the advanced computer vision models developed in our study together with the XAI model interpretation suggests that the high-performance computer vision models are reaching an upper bound of species classification performance on existing microscopy image datasets. Thus, increasing the amount of training data is expected to produce diminishing returns, with further improvements to classification performance requiring advances in experimental data collection and computational data processing to enhance species-distinguishing features, especially when expanding the models to multi-species classification tasks, as observed in other image-based deep learning studies^68^. This may be achieved using high-resolution single-cell imaging, staining methods, or computational feature enhancement. Additional XAI methods could interpret predictions on mixed-species cultures, such as Grad-CAM++^69^, which has better object localization and explanation of occurrences of multiple objects in a class compared to Grad-CAM^50^.

Each pathogenic yeast species considered in this study has a distinct size, shape, cell wall thickness, curvature, internal structure/organelle distribution, budding pattern, and optical properties^70^. This unique distribution of visual features that can be captured through bright-field microscopy, which our advanced computer vision models learned to make accurate species classification predictions. However, there are overlaps in the visual feature distributions (Figure 5, Supplementary Video 1, and Figure 7), which led to incorrect model predictions when the cell images do not contain sufficient distinguishing information. Other methods are capable of generating unique fingerprints of each species for accurate classification, including MALDI TOF MS by generating mass/charge ratio spectra^10^, Raman spectroscopic methods that detect vibrational signatures of molecules within the yeast cells^18^, using unique volatile organic compounds that are produced by each species^13^, and DNA sequencing^8^. However, we demonstrated that our models can successfully classify pathogenic yeast species based on their unique visual fingerprints captured by bright-field microscopy, while having a substantially lower resource cost compared to these other methods.

Our study highlights the power of advanced computer vision models to identify human fungal pathogens and XAI to elucidate the biologically and non-biologically relevant features responsible for model predictions. We anticipate that this approach of combining computer vision models with XAI will advance antimicrobial resistance research and lead to novel diagnostic methods to improve the treatment of patients with infectious diseases.

## 5 Acknowledgments

This research was funded by grants to D.A.C. from the Human Frontier Science Program (RGEC30/2024) and the Alberta Innovates Accelerating Innovations into Care - Concepts program (AICE-Concepts 597389). We thank Dr. Tanis Dingle and the Alberta Precision Laboratories-Public Health Laboratory for providing the *Candida* species isolates, Asst. Prof. Michael Manhart for suggesting the use of principal component analysis, and Prof. Randy Goebel for helpful discussions on explainable AI.

## 6. Author Contributions Statement

DAC, SAS, and JDG conceptualized the study. JDG carried out the computational work, data visualization, and analytics. SAS carried out the experimental work. SAS and JDG interpreted the results. JDG, SAS, and DAC wrote the manuscript. DAC acquired the funding and supervised the research.

## 7 Data Availability

The data that support the findings of this study are available upon reasonable, noncommercial request from the corresponding author.

