## Supplemental Information for "Peering Inside the Black Box: Explainable AI to Interpret Advanced Computer Vision Fungal Pathogen Prediction"

#### 1 Training, Validation, and Test Datasets

The training, validation, and test datasets are described in Supplementary Table 1. To show the standardization of the data collection, pixel distributions of the final training, validation, and test sets are shown in Supplementary Figures 1, 2, and 3, respectively. Each of these figures shows histograms of the average pixel value, standard deviation of the pixel values, and image size (before resizing for the models) for all images of each species in the dataset. Pixel values equal to zero (the black background) were ignored when calculating the average pixel values and standard deviations to highlight that the microscopy image collection was standardized across species. All images were cropped with a 1:1 square aspect ratio, so the image size histograms show only one dimension (e.g. 150 refers to a 150x150 image). These distributions show that all images were collected using the same microscope settings, resulting in nearly identical pixel distributions for all species in each dataset. The image sizes for each species are consistent across datasets but vary from species to species due to the different cell sizes.

#### 2 DenseNet121, InceptionV3, Vision Transformer, and Swin Transformer Test Results

The validation and test set performance metrics for the DenseNet121 model presented in the main text are shown in Supplementary Table 2a and 2b, respectively. Test set confusion matrices and performance metrics for the other three model architectures considered are shown in Supplementary Figure 4 and Supplementary Table 3, respectively. The similarity of the classification patterns indicated in the confusion matrices across each of the model architectures considered indicates that the models learned similar representations to reach optimal classification performance on the current data, adding further support to our hypotheses regarding the limitations of the current visual data described in the main text.

#### 3 DenseNet121 Results on Fully Automated Test Set Containing Outliers

We applied the DenseNet121 model described in the main text to the test set before the outliers were manually removed (Test Images P3 in Supplementary Table 1) to determine the impact the outliers have on the results when using fully automated data processing. This resulted in a slight decrease in overall accuracy to 86% compared to the manually cleaned test set (Supplementary Figure 5 shows the corresponding confusion matrix and Supplementary Table 4 shows the performance metrics). Comparing the confusion matrices and performance metrics of the uncleaned and manually cleaned test sets shows that all species other than *C. auris* and *P. krudiavzevii* (the only two species with non-negligible error rates) had little to no change in overall performance when the outliers were removed (*N. glabratus* had a 1% lower precision for the manually cleaned test set, and there was no change in the metrics of the other species). For the manually cleaned test set, *C. auris* had a 4% improvement in recall and a 1% increase in precision, while *P. krudiavzevii* had a 1% increase in recall and a 4% increase in precision. This suggests that many of the *C. auris* outliers were misidentified as *P. krudiavzevii*, which was confirmed by visually inspecting the predictions for the outliers. These results highlight the considerations that must be made when applying our methods in a fully automated setting where manual outlier removal may be infeasible.

#### 4 Comparison to Previous Work

To highlight the improvements we have made in image collection, data processing, model development and training to improve model generalization over previous work, we trained and evaluated models using only the 4 species previously investigated in Shankarnarayan & Charlebois, *Medical Mycology*, 2024[1] (*C. albicans*, *C. auris*, *N. glabratus*, and *C. haemulonii*), where an InceptionV3 model trained using transfer-learning (but not full model fine-tuning) had an overall test accuracy of 78%. After training the 4 species models as described in the main text using our new methods, each of the model architectures produced similar test set performance (Supplementary Figure 6a), with the Vision Transformer architecture producing the best results in this case with an overall accuracy of 92%. Supplementary Figure 6b shows the test set confusion matrix for the Vision Transformer model and Supplementary Figure 6c shows a comparison of species accuracies (recalls) between the Vision Transformer model and the InceptionV3 results published in Shankarnarayan & Charlebois, *Medical Mycology*, 2024[1]. Furthermore, our test set contained 3,827 cell images per species compared to 200 per species used in the previous work, providing additional support for our generalization improvement.

#### Supplementary Tables

| Yeast Species | Training Images | | | | Validation Images | | | | Test Images | | | | | |
|---|---|---|---|---|---|---|---|---|---|---|---|---|---|---|
| | Raw | P1 | P2 | P3 | Raw | P1 | P2 | P3 | Raw | P1 | P2 | P3* | P4 | P5 |
| *C. albicans* | 1,200 | 67,751 | 46,040 | 32,809 | 100 | 8,774 | 6,289 | 4,214 | 250 | 18,310 | 12,807 | 4,214 | 4,212 | 3,827 |
| *C. auris* | 1,000 | 69,462 | 32,809 | 32,809 | 300 | 59,261 | 19,781 | 4,214 | 100 | 13,390 | 6,720 | 4,214 | 3,827 | 3,827 |
| *N. glabratus* | 1,000 | 52,003 | 35,323 | 32,809 | 250 | 7,754 | 5,536 | 4,214 | 250 | 13,517 | 9,706 | 4,214 | 4,207 | 3,827 |
| *C. haemulonii* | 900 | 66,483 | 37,841 | 32,809 | 300 | 12,957 | 6,731 | 4,214 | 250 | 31,871 | 18,279 | 4,214 | 4,174 | 3,827 |
| *P. krudiavzevii* | 1,000 | 125,692 | 53,579 | 32,809 | 100 | 12,533 | 5,349 | 4,214 | 100 | 12,532 | 5,591 | 4,214 | 4,010 | 3,827 |
| *C. parapsilosis* | 1,000 | 135,421 | 84,816 | 32,809 | 150 | 6,536 | 4,214 | 4,214 | 100 | 27,119 | 17,348 | 4,214 | 4,174 | 3,827 |
| *C. tropicalis* | 1,000 | 99,352 | 67,914 | 32,809 | 100 | 7,234 | 4,963 | 4,214 | 100 | 11,168 | 8,115 | 4,214 | 4,189 | 3,827 |
| **Total** | 7,100 | 616,164 | 358,322 | 229,663 | 1,300 | 115,049 | 52,863 | 29,498 | 1,150 | 127,907 | 78,566 | 29,498 | 28,793 | 26,789 |

**Supplementary Table 1.** Training, validation, and test image datasets collected for the yeast species classification models. Individual cell images were cropped from raw bright-field microscope images of each species in several data processing (P) steps. Number of crops obtained after: applying segmentation and image processing (P1), using k-means to remove low-quality images (P2), randomly downsampling to match the species with the lowest number from P2 to maintain balanced datasets (P3), manually removing remaining outliers (P4), and downsampling to match the lowest number species after P4 (P5). *Test set P3 was downsampled to match the validation set size. The final datasets of individual cell images used for model training, validation, and testing are highlighted in grey.

**(a)**

| Species | Precision | Recall | $F_1$-Score |
|---|---|---|---|
| *C. albicans* | 0.83 | 0.92 | 0.87 |
| *C. auris* | 0.94 | 0.92 | 0.93 |
| *N. glabratus* | 0.91 | 0.98 | 0.94 |
| *C. haemulonii* | 0.79 | 0.77 | 0.78 |
| *P. krudiavzevii* | 0.91 | 0.90 | 0.90 |
| *C. parapsilosis* | 0.88 | 0.81 | 0.85 |
| *C. tropicalis* | 0.90 | 0.85 | 0.87 |
| **Averages** | 0.88 | 0.88 | 0.88 |

**(b)**

| Species | Precision | Recall | $F_1$-Score |
|---|---|---|---|
| *C. albicans* | 0.90 | 0.89 | 0.89 |
| *C. auris* | 0.94 | 0.90 | 0.92 |
| *N. glabratus* | 0.92 | 0.85 | 0.88 |
| *C. haemulonii* | 0.90 | 0.76 | 0.83 |
| *P. krudiavzevii* | 0.88 | 0.90 | 0.89 |
| *C. parapsilosis* | 0.73 | 0.97 | 0.83 |
| *C. tropicalis* | 0.88 | 0.82 | 0.85 |
| **Averages** | 0.88 | 0.87 | 0.87 |

**Supplementary Table 2.** Validation and test performance metrics for the DenseNet121 model. **(a)** Validation set performance metrics (overall validation accuracy of 88%). Each species had 4214 validation samples. **(b)** Test set performance metrics (overall test accuracy of 87%). Each species had 3827 test samples.

**(a)**

| Species | Precision | Recall | $F_1$-Score |
|---|---|---|---|
| *C. albicans* | 0.84 | 0.91 | 0.87 |
| *C. auris* | 0.94 | 0.94 | 0.94 |
| *N. glabratus* | 0.92 | 0.76 | 0.83 |
| *C. haemulonii* | 0.90 | 0.75 | 0.82 |
| *P. krudiavzevii* | 0.87 | 0.89 | 0.88 |
| *C. parapsilosis* | 0.71 | 0.96 | 0.82 |
| *C. tropicalis* | 0.89 | 0.78 | 0.83 |
| **Averages** | 0.87 | 0.86 | 0.86 |

**(b)**

| Species | Precision | Recall | $F_1$-Score |
|---|---|---|---|
| *C. albicans* | 0.85 | 0.90 | 0.88 |
| *C. auris* | 0.93 | 0.93 | 0.93 |
| *N. glabratus* | 0.96 | 0.78 | 0.86 |
| *C. haemulonii* | 0.90 | 0.75 | 0.82 |
| *P. krudiavzevii* | 0.82 | 0.90 | 0.86 |
| *C. parapsilosis* | 0.73 | 0.96 | 0.83 |
| *C. tropicalis* | 0.87 | 0.77 | 0.82 |
| **Averages** | 0.87 | 0.86 | 0.86 |

**(c)**

| Species | Precision | Recall | $F_1$-Score |
|---|---|---|---|
| *C. albicans* | 0.88 | 0.90 | 0.89 |
| *C. auris* | 0.95 | 0.94 | 0.94 |
| *N. glabratus* | 0.93 | 0.78 | 0.85 |
| *C. haemulonii* | 0.92 | 0.68 | 0.78 |
| *P. krudiavzevii* | 0.85 | 0.90 | 0.88 |
| *C. parapsilosis* | 0.69 | 0.98 | 0.81 |
| *C. tropicalis* | 0.88 | 0.83 | 0.85 |
| **Averages** | 0.87 | 0.86 | 0.86 |

**Supplementary Table 3.** Test set performance metrics for the additional model architectures considered. **(a)** InceptionV3. **(b)** Vision Transformer (Base 16). **(c)** Swin Transformer (Tiny).

| Species | Precision | Recall | $F_1$-Score |
|---|---|---|---|
| *C. albicans* | 0.90 | 0.89 | 0.89 |
| *C. auris* | 0.93 | 0.86 | 0.90 |
| *N. glabratus* | 0.93 | 0.85 | 0.89 |
| *C. haemulonii* | 0.90 | 0.76 | 0.82 |
| *P. krudiavzevii* | 0.84 | 0.89 | 0.87 |
| *C. parapsilosis* | 0.73 | 0.97 | 0.83 |
| *C. tropicalis* | 0.88 | 0.82 | 0.85 |
| **Averages** | 0.87 | 0.86 | 0.86 |

**Supplementary Table 4.** Performance metrics for the DenseNet121 model on the test set without the outliers manually removed (overall accuracy of 86%). Each species had 4214 test samples, with 2.4% overall being outliers.

### Supplementary Figures

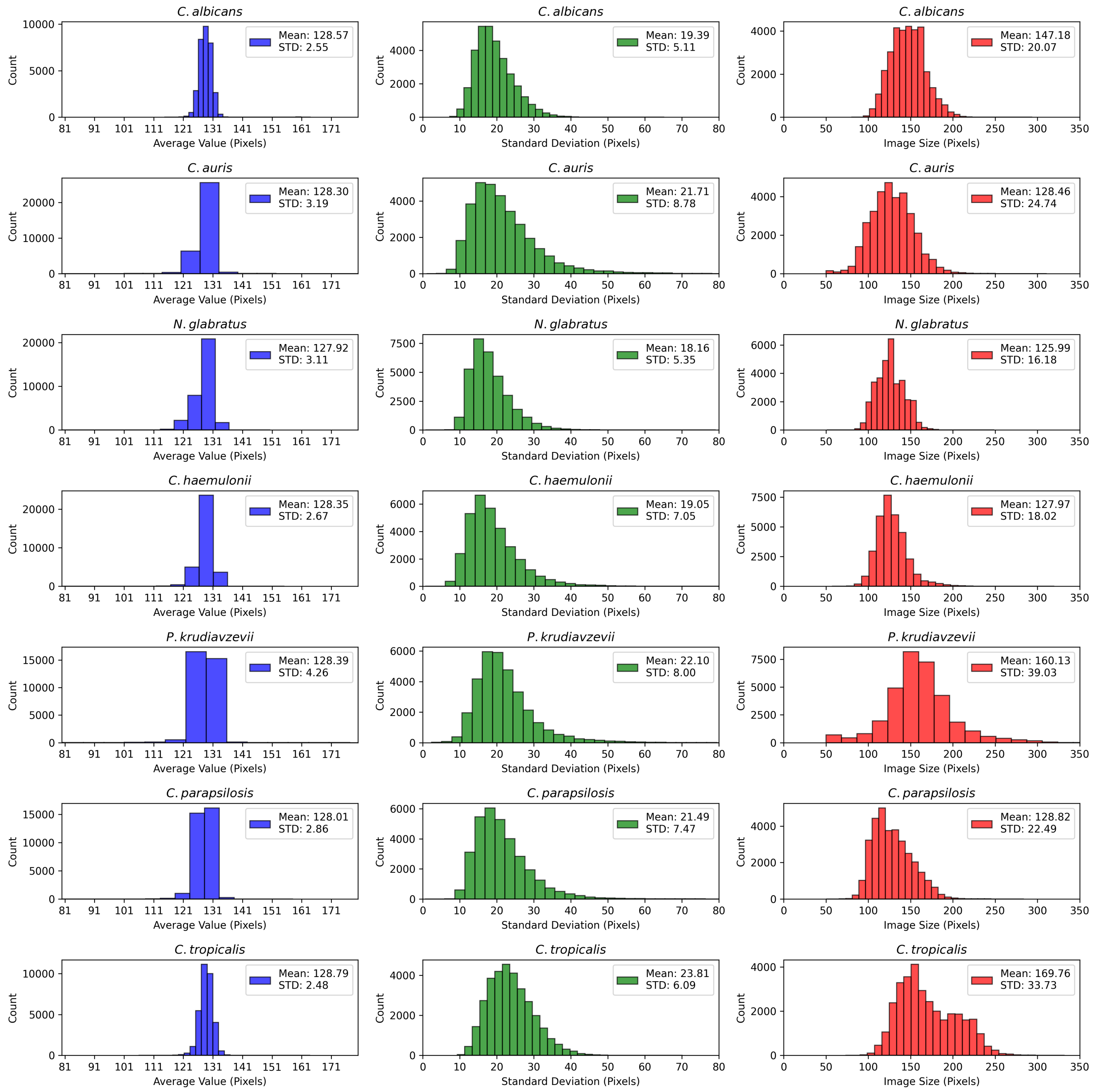


**Supplementary Figure 1.** Training dataset pixel distributions and image sizes for each species.

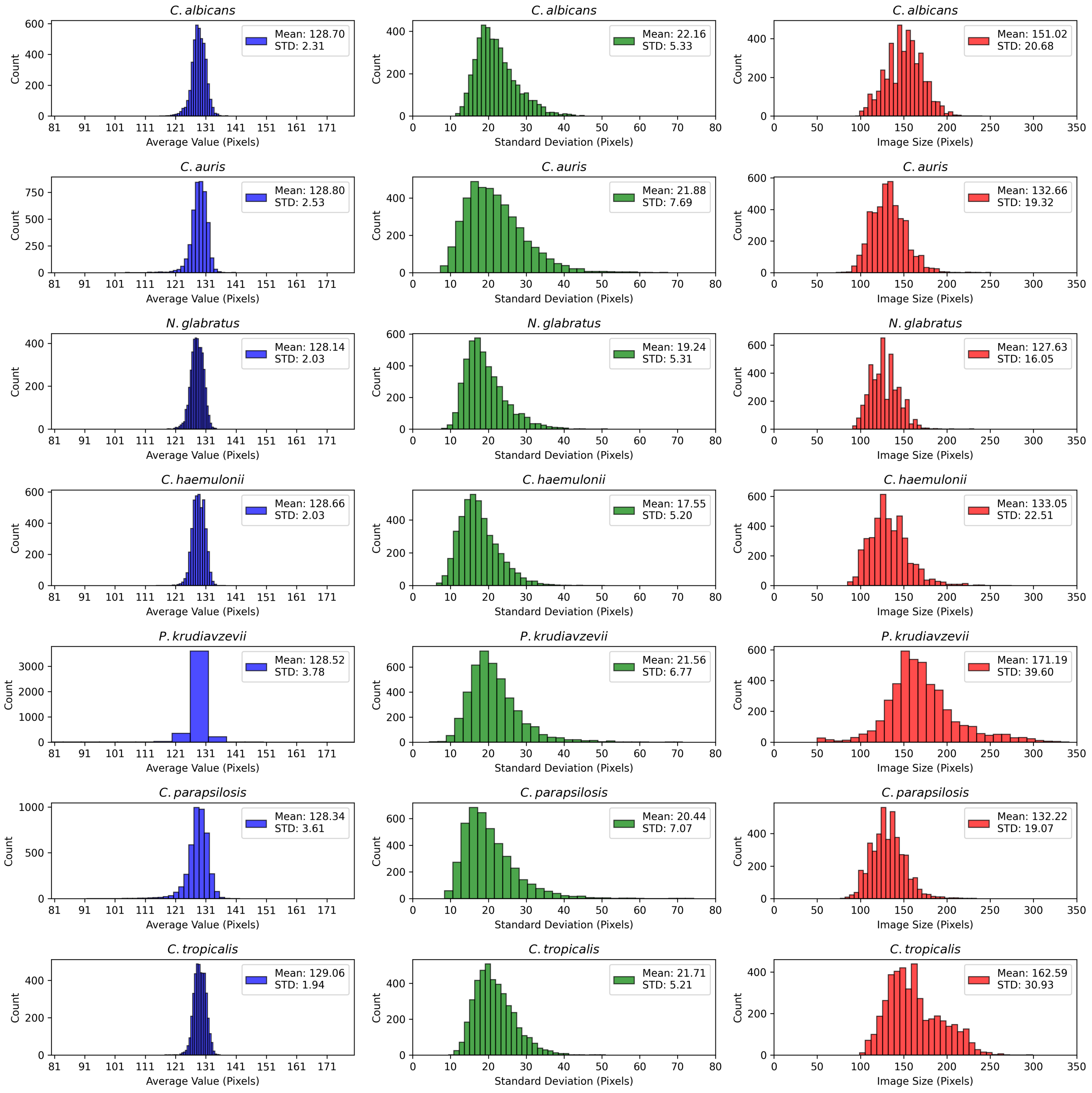


**Supplementary Figure 2.** Validation dataset pixel distributions and image sizes for each species.

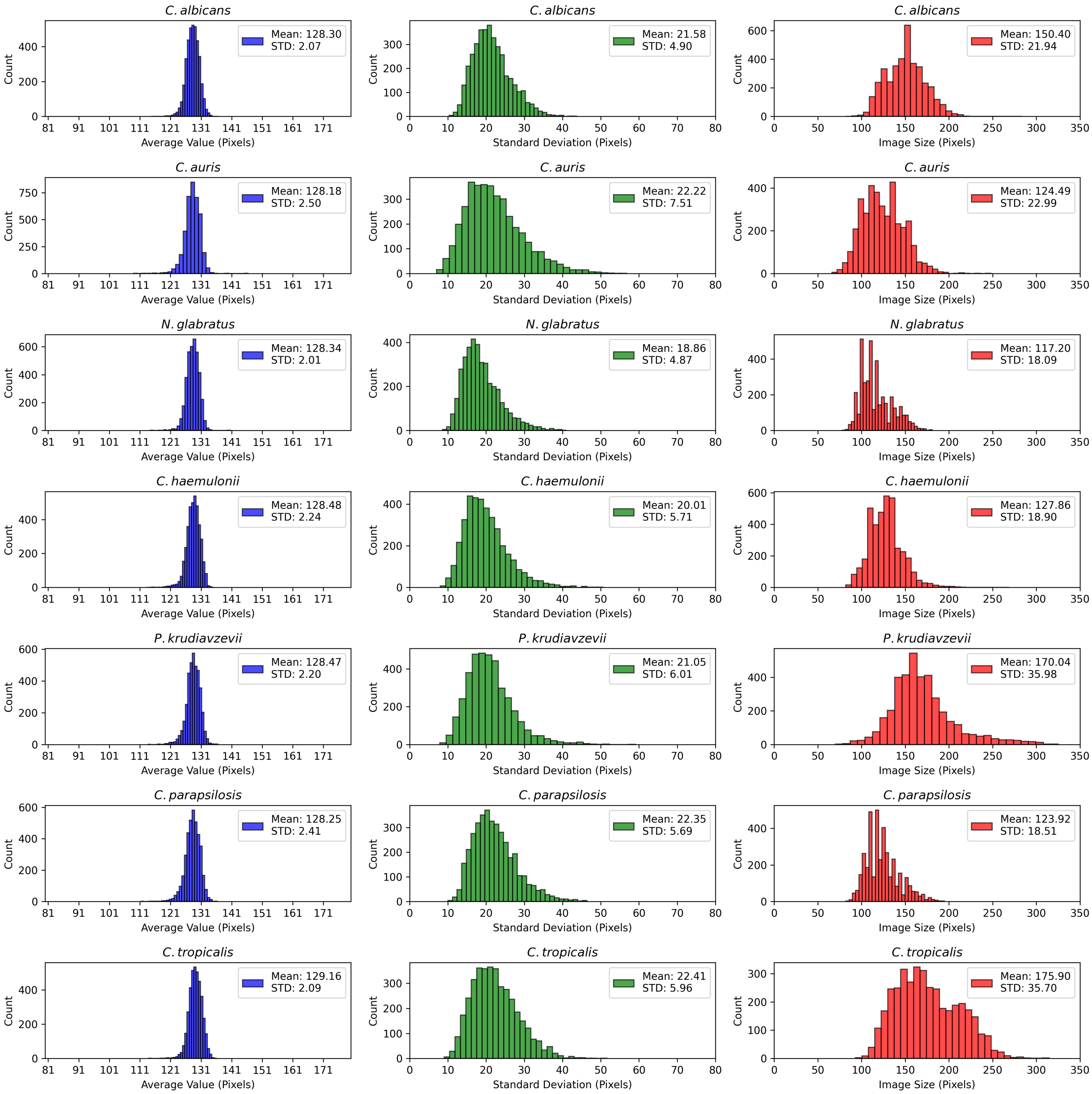


**Supplementary Figure 3.** Test dataset pixel distributions and image sizes for each species.

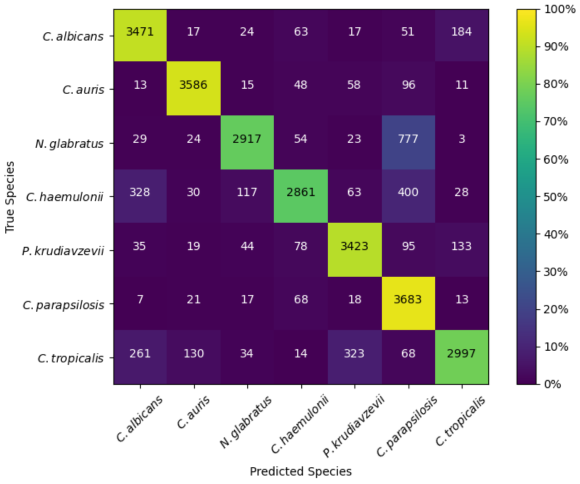


(a)

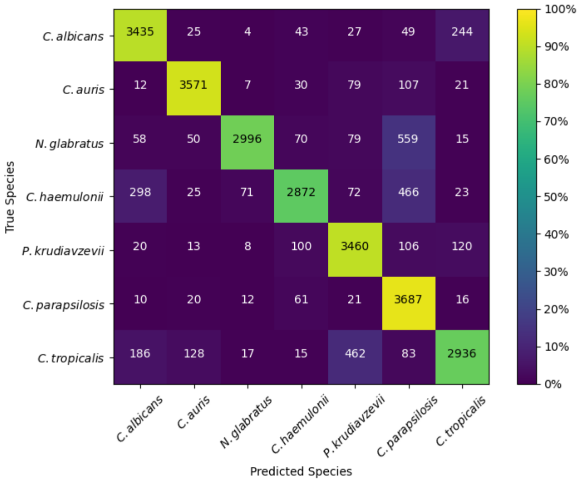


(b)

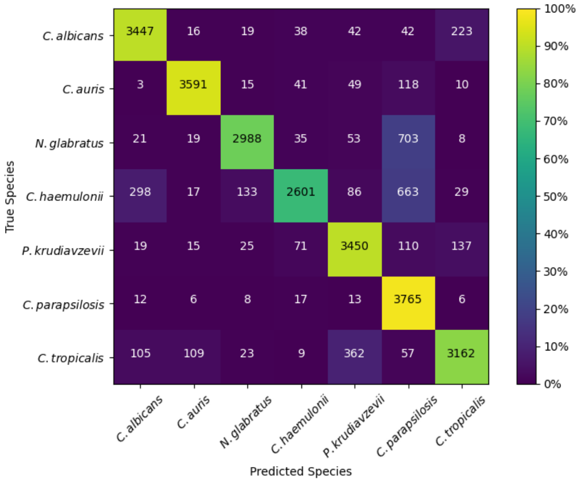


(c)

**Supplementary Figure 4.** Test set confusion matrices for the additional model architectures considered. **(a)** InceptionV3. **(b)** Vision Transformer (Base 16). **(c)** Swin Transformer (Tiny).

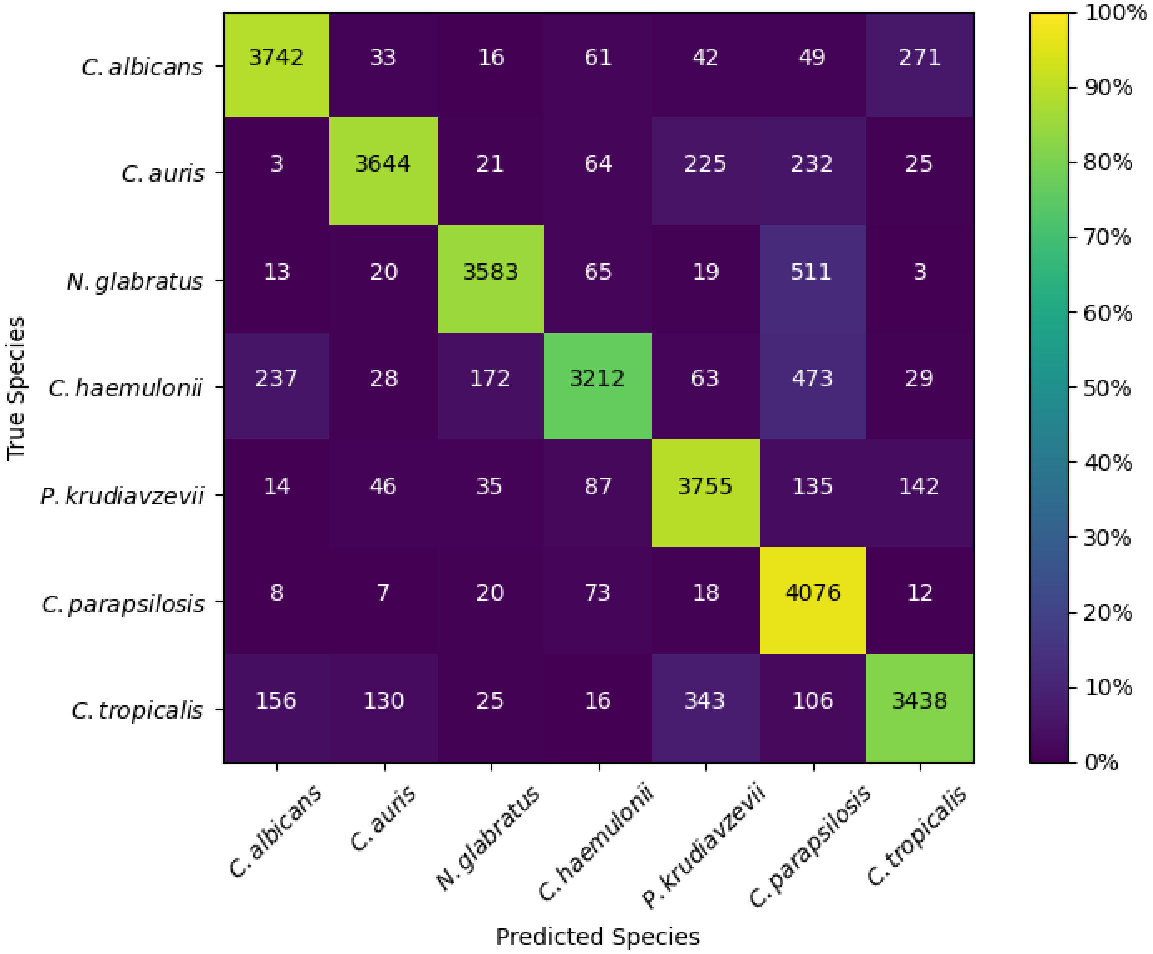


**Supplementary Figure 5.** Confusion matrix for the DenseNet121 model on the test set before the outliers were manually removed. The heatmap shows the percentage of samples in each cell relative to the dataset size of each class (4214, where 2.4% overall are outliers).

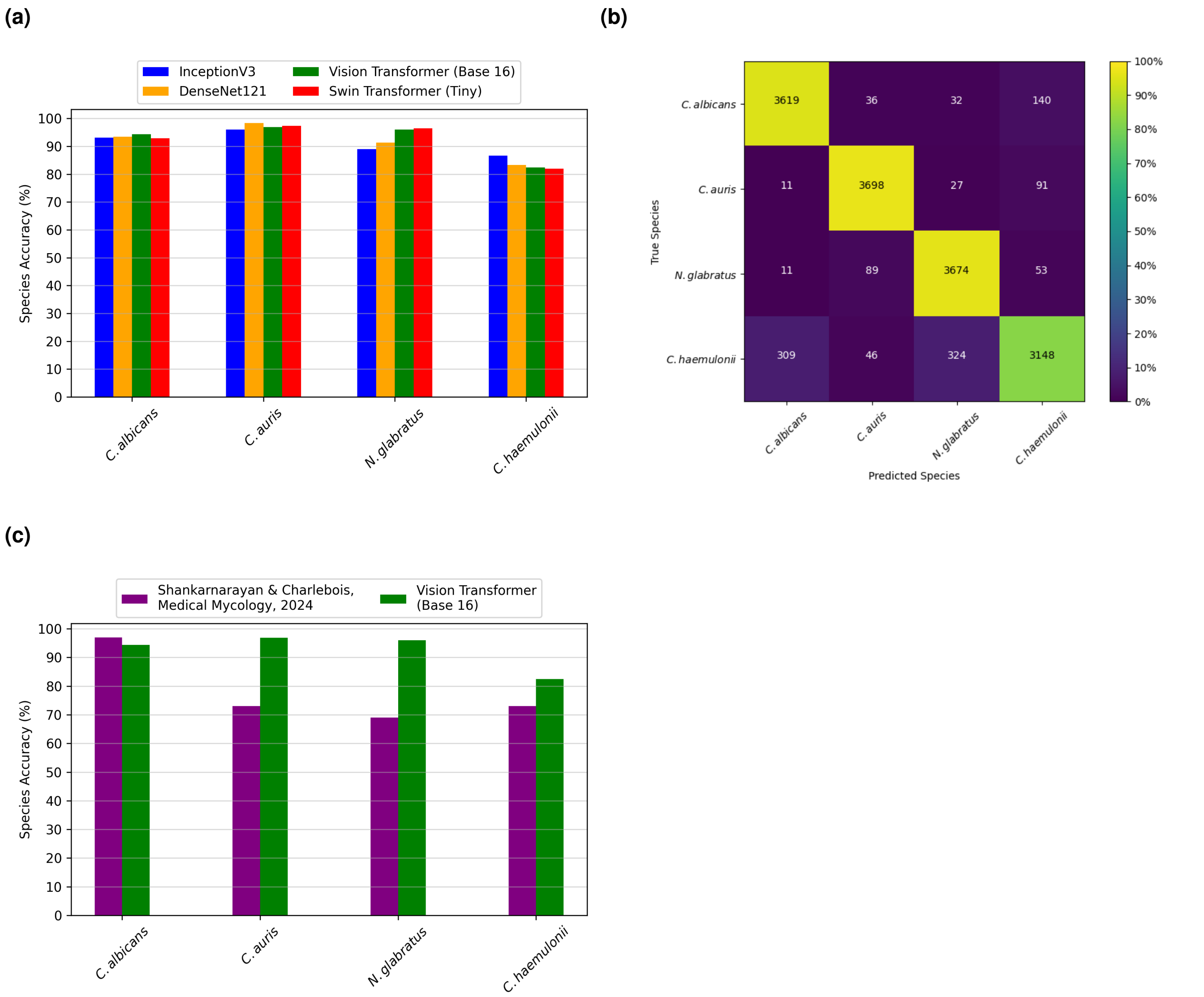


**Supplementary Figure 6.** Results using the 4 species (*C. albicans*, *C. auris*, *N. glabratus*, *C. haemulonii*) previously investigated in Shankarnarayan & Charlebois, *Medical Mycology*, 2024[1]. **(a)** Species accuracy (recall) comparison between the model architectures for the 4 species. **(b)** Confusion matrix when applying the 4-species Vision Transformer model to the test set (overall accuracy 92%). The heatmap shows the percentage of samples in each cell relative to the dataset size of each class (3827). **(b)** Species test accuracy (recall) comparison between our 4-species Vision Transformer model and the InceptionV3-based model previously published by Shankarnarayan & Charlebois[1].
